# Short repeats drive mitochondrial genome expansion and record hybridization in *Camellia*

**DOI:** 10.64898/2026.03.16.711854

**Authors:** Fen Zhang, Li-zhi Gao

**Author notes:** Corresponding author: Li-zhi Gao.

## Abstract

Plant mitochondrial genomes are paradoxical: structurally dynamic yet functionally conserved. In hybrid-prone lineages like *Camellia* (Theaceae), the forces shaping mitogenome evolution remain unclear. Here, we assemble mitogenomes of eight *Camellia* species and an outgroup. We show that short repeats (<100 bp) drive genome expansion (PGLS: *R*² = 0.97, *P* = 6.3 × 10⁻⁶), overturning the long-repeat paradigm. Multichromosomal architectures and extensive horizontal transfer (up to 25.5% plastid DNA) characterize hybrid lineages. Stark phylogenomic conflicts among mitochondrial, chloroplast and nuclear genomes expose genome-wide hybridization signatures. Mitochondria thus act as recorders of historical gene flow, capturing repeat explosions triggered by introgression. This work redefines plant mitogenome evolution and provides a framework to resolve taxonomic complexities in rapidly radiating angiosperms.

## Introduction

Mitochondrial genomes in flowering plants are evolutionary paradoxes, conserved in core protein-coding genes yet wildly divergent in structure and size, shaped by rampant recombination, horizontal gene transfer (HGT), and repeat-driven expansion^1–4^. These enigmatic genomes, often spanning hundreds of kilobases to megabases, challenge the streamlined mitochondrial archetype of animals and fungi^5–7^. While their role in energy production is well-established, their contributions to speciation and hybridization, particularly in rapidly radiating plant groups, remain understudied. Recent work in *Arabidopsis* and *Mimulus* has revealed mitochondrial-nuclear incompatibilities that drive hybrid breakdown and ecological adaptation^8–10^, yet systematic comparisons of mitochondrial diversity in taxonomically contentious lineages are lacking. This gap is especially pronounced in genera like *Camellia*, where hybridization and polyploidy blur species boundaries, complicating phylogenetic reconstruction and obscuring evolutionary mechanisms^11–14^.

The genus *Camellia*, home to the tea plant (*C. sinensis*) and over 100 species of ecological and economic importance, epitomizes the challenges of studying mitochondrial evolution in hybrid-rich systems^15,16^. Frequent interspecific hybridization, recurrent polyploidization, and incomplete lineage sorting have rendered *Camellia* a notorious “taxonomic labyrinth,” with nuclear and chloroplast phylogenies often conflicting^17,18^. Mitochondrial genomes, maternally inherited and shielded from recombination in most angiosperms, could provide a stable phylogenetic signal to resolve these conflicts^19,20^. However, mitochondrial genome data have been absent from *Camellia* systematics, reflecting broader neglect of cytoplasmic genomes in plant radiations shaped by reticulate evolution. This oversight persists despite evidence from *Quercus* and *Rhododendron*, where mitochondrial phylogenies clarify relationships obscured by nuclear introgression^21–23^.

Hybridization and polyploidy, two pillars of angiosperm diversification, leave distinct genomic imprints. In nuclear genomes, these include homoeologous exchanges and subgenome dominance; in chloroplasts, biparental inheritance often fuels cytonuclear discordance^24,25^. Mitochondrial genomes, however, respond uniquely, their low recombination rates and susceptibility to HGT may amplify structural variation during genomic upheavals. Polyploid wheat, for example, exhibits mitochondrial genome expansion via retrotransposon proliferation^26^, while hybrid *Helianthus* species show elevated plastid DNA integration into mitochondria^27^. In *Camellia*, where hybridization and polyploidy are pervasive^11–14^, mitochondrial genomes likely harbor signatures of these processes, yet their structural and evolutionary dynamics remain uncharacterized.

Here, we resolve this knowledge gap through comparative analysis of complete mitochondrial genomes across eight representative *Camellia* species and the outgroup *Actinidia eriantha* through PacBio Sequel platform. Our study reveals that *Camellia* mitochondrial genomes are dominated by short tandem repeats (STRs) (<100 bp), which drive ∼92.6–94.9% of genome size variation (*R*²=0.9729, *P*=6.293×10⁻⁶), a stark departure from models emphasizing large repeats in plants like *Citrullus lanatus* and *Cucurbita pepo*^28^. Lineage-specific blocks of plastid-derived DNA, occupying up to 25.5% of mitochondrial content, correlate with historical hybridization events inferred from nuclear data^29^, positioning mitochondrial genomes as archives of past gene flow. Strikingly, mitochondrial phylogenies resolve previously intractable relationships, such as the sister grouping of *C. sinensis* varieties CSSBY and CSSLJ43 despite chloroplast discordance. Species with strong hybridization signatures (e.g., *C. reticulata*, CRE) exhibits mitochondrial expansion (1.2× sister species) via retrotransposon-like repeats. These findings redefine mitochondrial roles in plant speciation. By demonstrating that mitochondrial genomes capture hybridization through structural innovation and foreign DNA acquisition, our work establishes *Camellia* as a model for studying cytoplasmic evolution in taxonomically complex radiations. The hyper-permissiveness of *Camellia* mitochondria to plastid DNA, exceeding levels in *Amborella*^30^, suggests HGT may mitigate cytoplasmic incompatibilities during hybridization, a hypothesis testable in experimental crosses. Moreover, the dominance of short repeats over large duplications in genome expansion challenges traditional models, urging reevaluation of repeat classification in plant mitochondrial studies. Beyond *Camellia*, this work provides a framework to resolve phylogenetic conflicts in hybrid-rich groups like *Rhododendron* and *Quercus*, while offering tools to engineer cytoplasmic compatibility in crops, an imperative as climate change reshapes agricultural landscapes^21–23^.

## Results

### Multichromosomal Architectures and Genome-Wide Structural Diversity

The mitochondrial genomes of five *Camellia* species and *Actinidia eriantha* as outgroup exhibited striking structural heterogeneity, ranging from single circular chromosomes to fragmented multichromosomal configurations (**Fig. 1**, **Table 1**). *C. lanceoleosa* (CLA) harbored the largest mitogenome (1,000,198 bp), while *C. reticulata* (CRE) possessed the smallest (788,876 bp), reflecting a 1.27-fold size variation within the genus (**Fig. 1**, **Table 1**). Crucially, two species, *C. sinensis* var. *sinensis* cv. Longjing43 (CSSLJ43) and *C. taliensis* (CTA), displayed fragmented mitogenomes partitioned into two circular chromosomes (Chr1: 767,375 bp and Chr2: 124,086 bp for CSSLJ43; Chr1: 862,525 bp and Chr2: 126,090 bp for CTA) (**Fig. 1**, **Table 1**). After remapping the reads back to the final assembly, we observed even read distribution across two chromosomes, suggesting the quality of genome assemblies (**Supplementary Fig. 1A**).

**Fig. 1.**
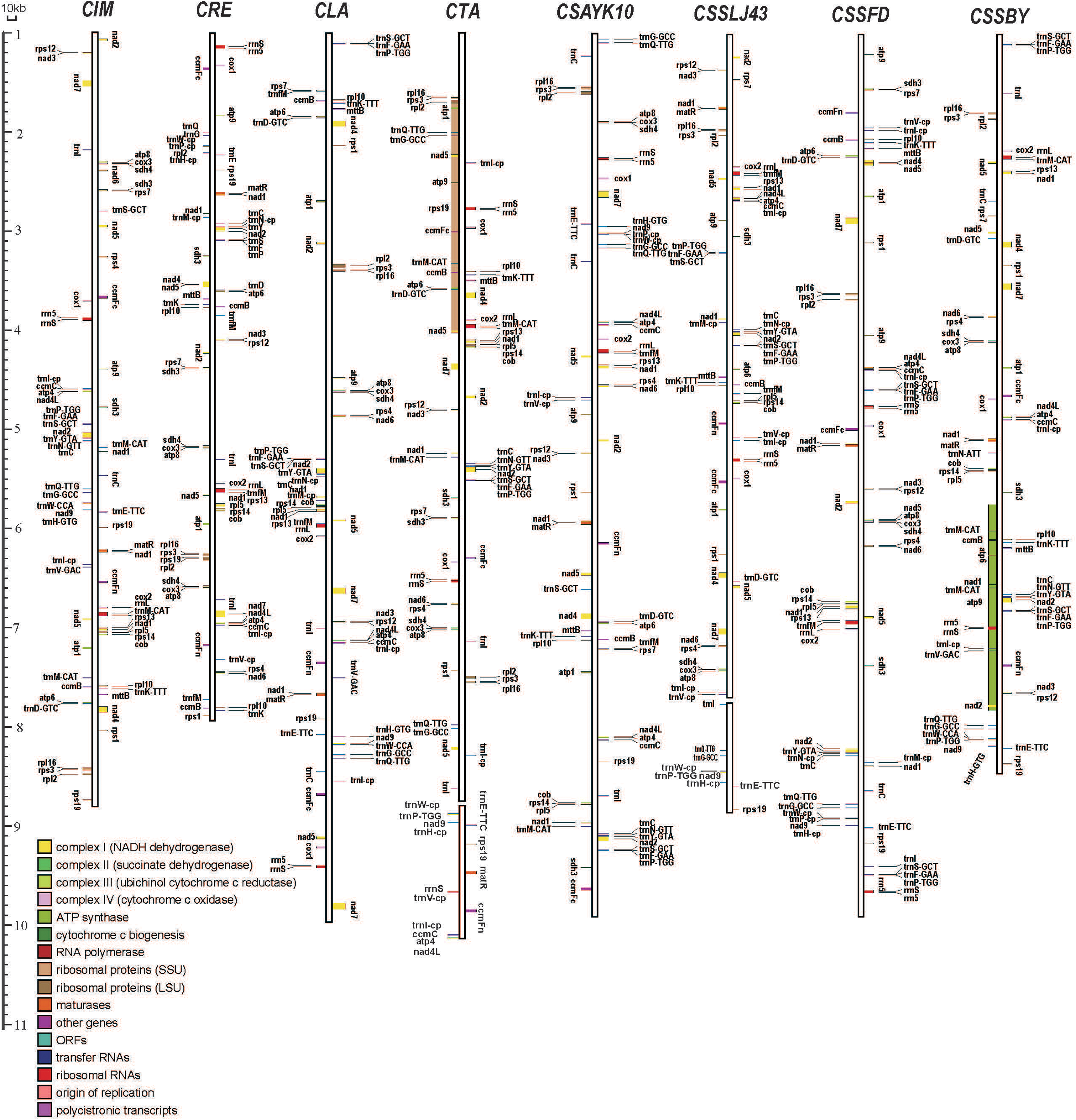
Structural organization of *Camellia* mitochondrial genomes. Linear representation of eight *Camellia* mitogenomes and the outgroup *Actinidia eriantha*. Genes on the left and right strands are transcribed clockwise and counterclockwise, respectively. Functional categories are color-coded. Chloroplast-derived sequences are marked with “-cp” suffixes. Genome assemblies were validated by split-read mapping (**Supplementary** Fig. 1). Scale bar: 10 kb. Chromosomal configurations (single-circular vs. multichromosomal) are indicated for *C. sinensis* var. *sinensis* LJ43 (CSSLJ43) and *C. taliensis* (CT). Gene order disruptions are highlighted at repeat junctions.

**Table 1.** Summary of features in the eight *Camellia* plant mitogenomes. SNP/Indel density and plastid-derived DNA indicate hybridization history in all species.

| Feature | CSSBY | CSSFD | CSSLJ43 | CSAYK10 | CTA | CLA | CRE | CIM | AER |
| --- | --- | --- | --- | --- | --- | --- | --- | --- | --- |
| Genome size (bp) | 840,303 | 987,158 | Chr1:767,375<br>Chr2:124,086 | 984,663 | Chr1:862,525<br>Chr2:126,090 | 1,000,198 | 788,876 | 887,673 | 816,746 |
| Number of chromosomes | 1 | 1 | 2 | 1 | 2 | 1 | 1 | 1 | 1 |
| GC content (%) | 45.78 | 45.61 | Chr1:45.59<br>Chr2:46.04 | 45.60 | Chr1:45.73<br>Chr2:45.78 | 45.72 | 46.01 | 45.72 | 46.17 |
| Number of protein genes | 38 | 40 | 38 | 42 | 46 | 38 | 45 | 40 | 42 |
| Number of tRNA genes | 26 | 24 | 26 | 26 | 25 | 24 | 23 | 22 | 18 |
| Number of rRNA genes | 3 | 5 | 3 | 3 | 6 | 3 | 3 | 3 | 4 |
| Total genes | 67 | 69 | 67 | 71 | 77 | 65 | 71 | 65 | 64 |
| Number of SSRs | 651 | 766 | 679 | 743 | 736 | 757 | 620 | 692 | 539 |
| Length of repeats (bp) | 21,894 | 245,392 | 117,118 | 248,817 | 479,439 | 176,922 | 176,922 | 70,783 | 104,740 |
| Percentage of repeats (%) | 2.61 | 24.86 | 13.14 | 25.27 | 48.5 | 17.69 | 22.43 | 7.97 | 12.82 |
| Longest repeat (bp) | 395 | 34,463 | 22,564 | 30,713 | 50,026 | 60,766 | 26,692 | 11,988 | 10,504 |
| Tandem Repeats | 28 | 46 | 39 | 36 | 46 | 41 | 32 | 31 | 23 |
| Number of RNA-editing | 518 | 514 | 524 | 527 | 505 | 523 | 500 | 522 | 517 |
| Number of SNP |  | 605 | 722 | 783 | 766 | 1,068 | 622 | 818 | 5,157 |
| Number of Indel |  | 545 | 551 | 669 | 656 | 717 | 622 | 607 | 1,369 |
| Length of plastid-derived sequences (bp) | 26,430 | 30,890 | 40,115 | 25,662 | 32,105 | 27,016 | 15,238 | 27,011 | 11,824 |
| Length of nuclear-shared sequences (bp) | 2,323,032 | 2,536,043 | 2,407,094 | 2,597,562 | 2,616,563 | 2,572,908 | 3,877,865 | 2,489,068 | 623,528 |
| Number of gene acquired by HGT | 57 | 60 | 59 | 60 | 58 | 57 | 59 | 58 | 57 |

Gene order comparisons revealed extreme collinearity disruption among *Camellia* mitogenomes. Synteny analysis using CSSBY as a reference showed that no collinear blocks exceeded 50 kb in length, with pairwise alignment identities dropping below 74% in noncoding regions (**Fig. 2A**). For instance, the *nad5-cox1* cluster, conserved in most angiosperms^31^, was fragmented into three segments in CTA, flanked by 34.5 kb of repeats (**Fig. 2B**). Structural variation density plots highlighted hotspots of SNPs and indels near repeat junctions (**Fig. 2C**, **Supplementary Table 1**), implicating recombination as a driver of gene order scrambling. Notably, CSSFD and CTA, despite similar genome sizes (987,158 bp vs. 988,615 bp), shared only 42% syntenic regions, underscoring rampant rearrangement (**Supplementary Fig. 2**).

**Fig. 2.**
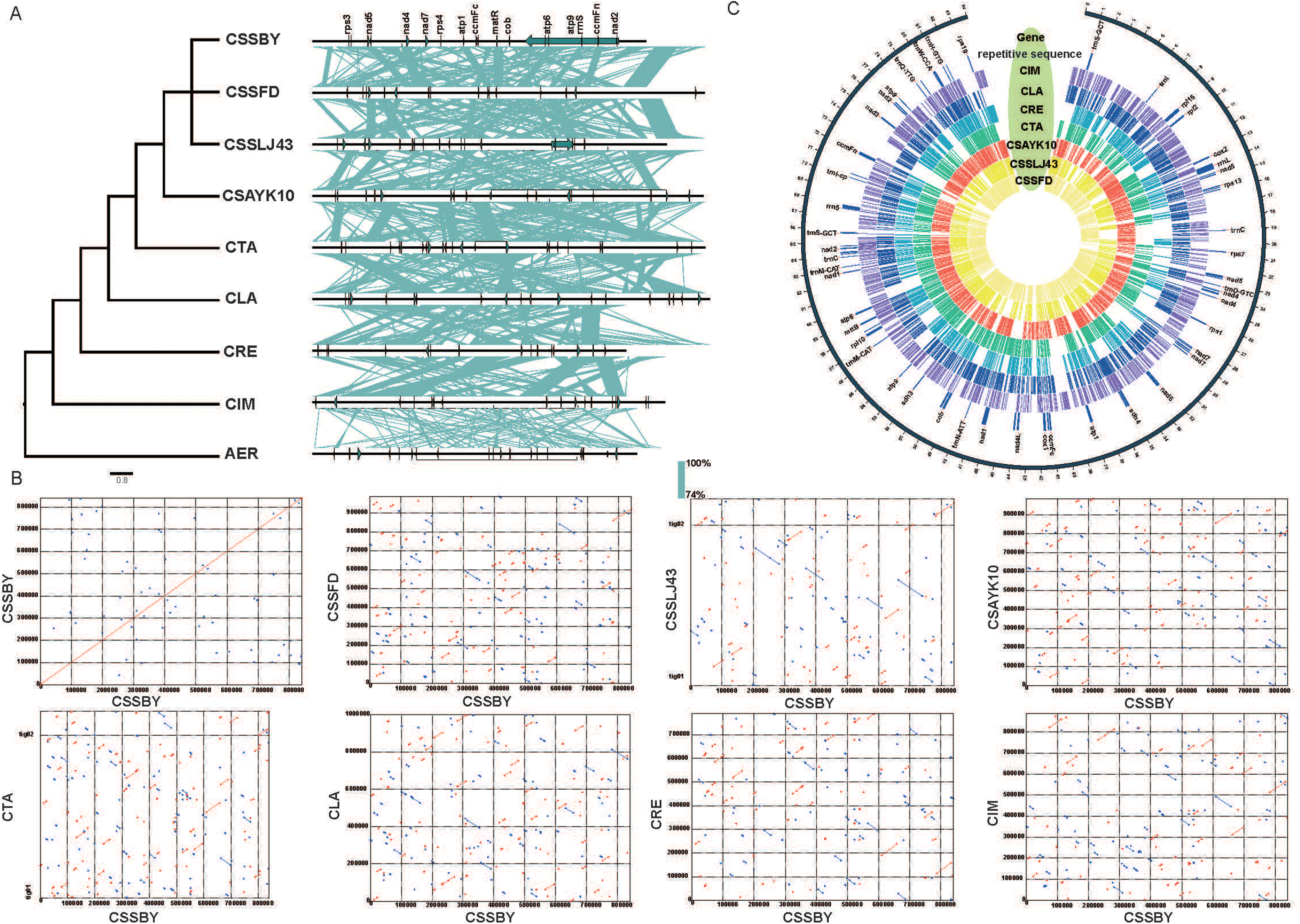
Structural variation and collinearity of *Camellia* mitogenomes. (A) Synteny network based on phylogenetic relationships. (B) Pairwise alignments using CSSBY as reference. Color intensity reflects sequence identity (74-100%). (C) SNP/indel density (variants/kb) across CSSBY mitogenome. Hotspots near repeat junctions (blue bars) suggest recombination-driven rearrangements. Intergenic regions show elevated polymorphism (4.2× vs. coding regions). Genome coordinates (Mb) and key genes are labeled.

### Repeat-Driven Genome Expansion: Short Repeats Dominate in Ericales

Comparative repeat analysis resolved a near-perfect correlation between genome size and total repeat number (PGLS: *R*² = 0.9729, *P* = 6.29×10⁻⁶ (**Fig. 3, Supplementary Fig. 3, Supplementary Table 2-5**). However, contrary to findings in *Monsonia*, *Citrullus lanatus* and *Cucurbita pepo* ^25,28^(short repeats <500 bp) dominated *Camellia* mitogenome expansion. Repeats <100 bp accounted for ∼92.6–94.9% of all repetitive elements (**Fig. 3A, Supplementary Table 3**), contributing ∼2–48% of total repeat length in most species (**Fig. 3B**). Strikingly, CTA’s mitogenome contained 479,439 bp of repeats (∼48.5% of total length), with ∼95.3% of repeat length attributed to sequences >1 kb (**Fig. 3B, Supplementary Table 3**), suggesting lineage-specific expansion mechanisms.

**Fig. 3.**
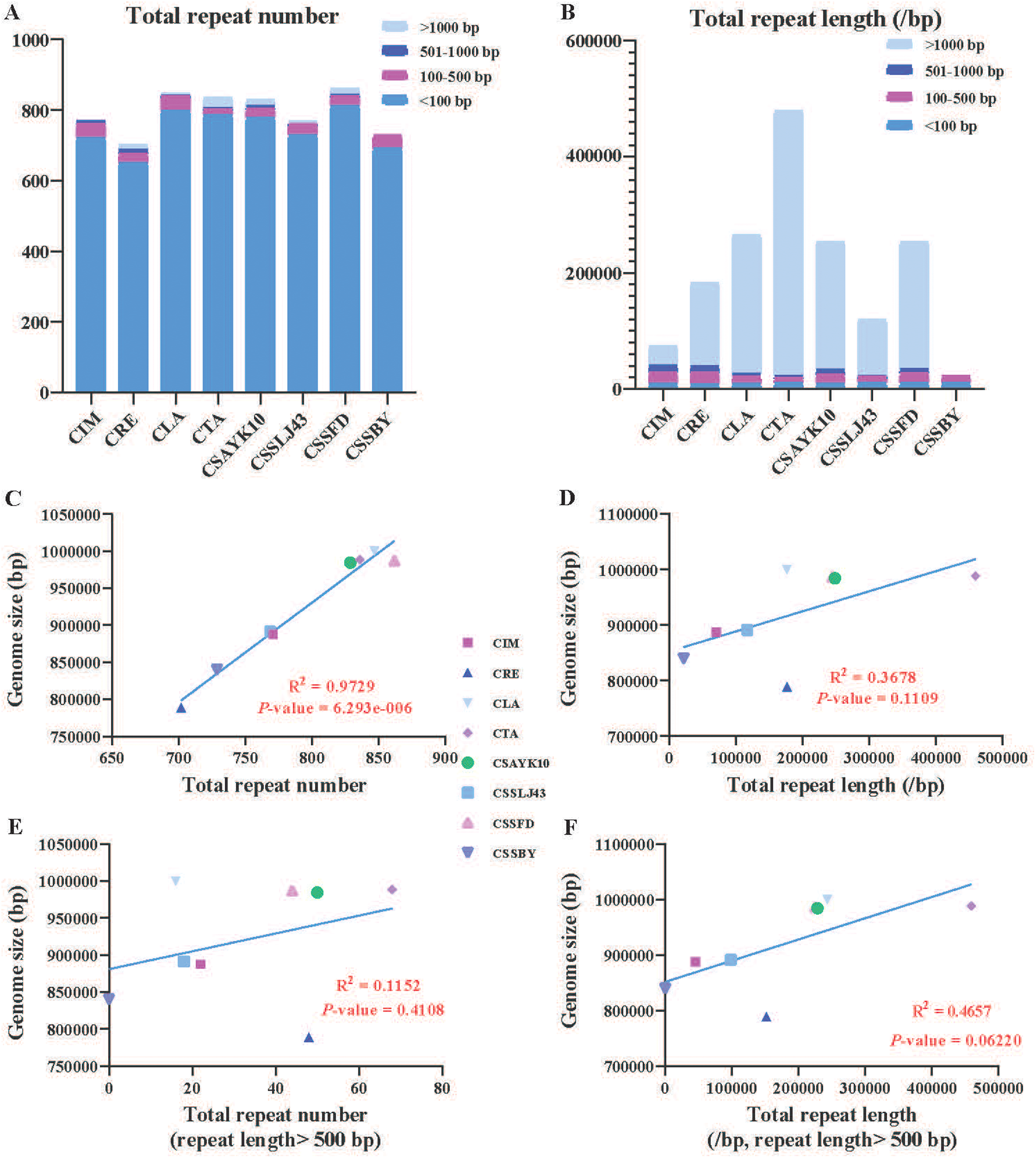
Repeat-driven expansion of *Camellia* mitochondrial genomes. (A) Total repeat counts classified by size: <100 bp, 100-500 bp, 500-1,000 bp, >1,000 bp. (B) Contribution of repeat length to genome size. (C) Genome size-repeat correlation across hybridizing diploids. Phylogenetic generalized least squares (PGLS) regression showing genome size correlates with total repeat number (R² = 0.9729, *P* = 6.293×10⁻⁶). (D-F) Subgroup analyses for repeats ≤500 bp. Data points represent individual species (n = 8); shaded areas indicate 95% confidence intervals.

SSR and tandem repeat (TR) profiling further highlighted genomic dynamism. CSSFD harbored the highest SSR density (766 SSRs), primarily dimers (348) and monomers (238), while CRE had the lowest (620 SSRs) (**Table 1, Supplementary Table 2**). TRs ≥6 bp with ≥70% identity ranged from 28 (CSSBY) to 46 (CSSFD, CTA) per genome (**Supplementary Table 4**). Long repeats (>50 bp) occupied up to 310 kb in CTA (**Supplementary Table 5**), often flanking structural breakpoints (**Fig. 2C**). These findings overturn the prevailing model that long repeats drive plant mitogenome inflation^25^, establishing short repeats as key players in Ericales.

### Horizontal Gene Transfer (HGT), Phylogenomic Discordance and Structural Diversity

Chloroplast-derived sequences (NUPTs) occupied ∼9.7–25.5% of mitochondrial content, with the *mtcp15* fragment uniquely shared among *C*.

*sinensis* var. *sinensis* cultivars (CSSBY, CSSFD, CSSLJ43; **Fig. 4A**). CSSLJ43 harbored the largest NUPT segment (40,115 bp), including a 22.3 kb inversion spanning the chloroplast inverted repeat (IR) region (**Fig. 4A**). Phylogenetically restricted transfers were identified: the *mtcp15* fragment, derived from the chloroplast *trnH-GUG* locus, was present only in three *C. sinensis* var. *sinensis* cultivars (CSSBY, CSSFD, CSSLJ43), suggesting post-divergence transfer after the *sinensis-assamica* split (**Fig. 4A**).

**Figure 4.**
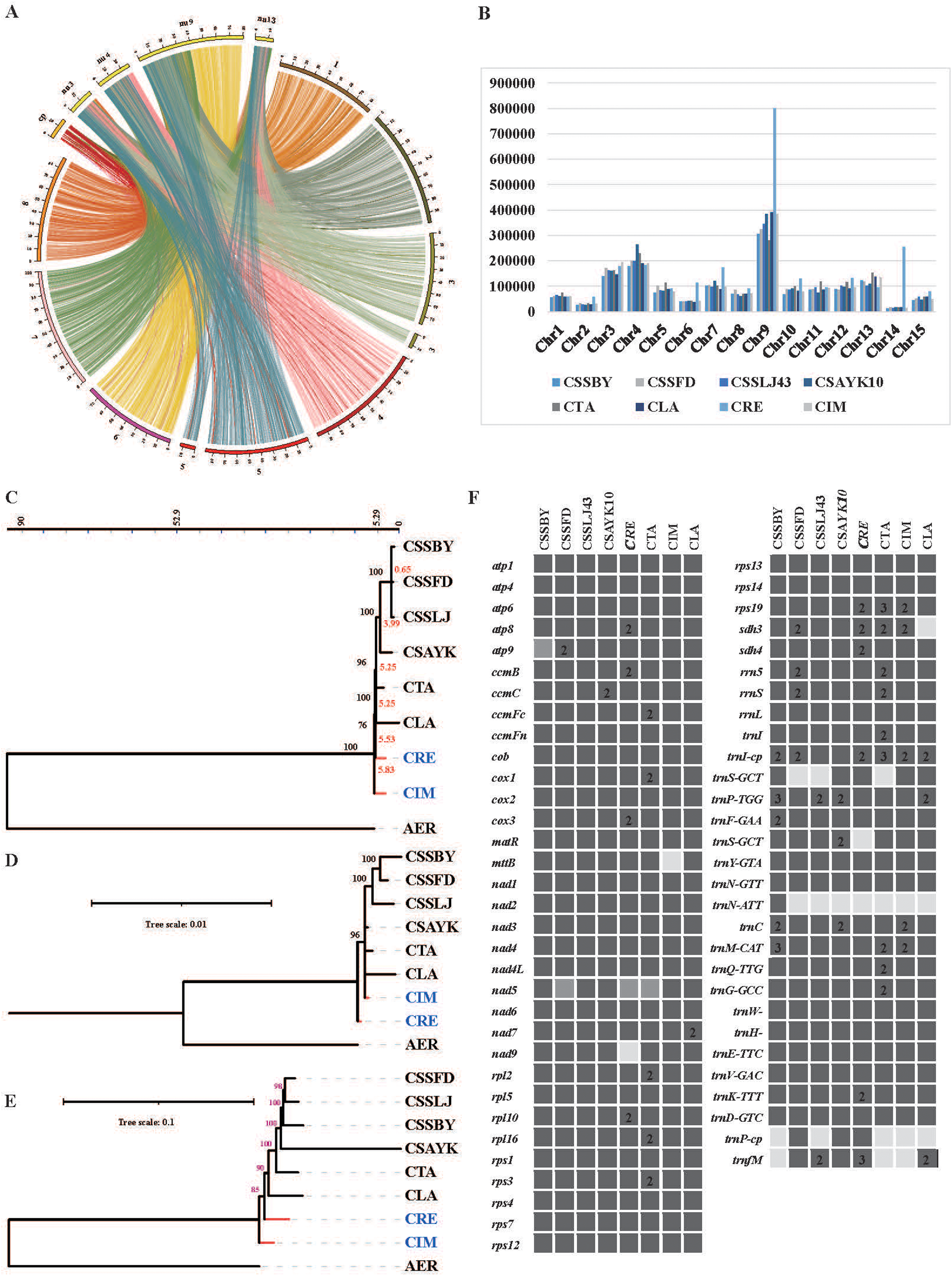
Mitochondrial genomes capture hybridization history through HGT, phylogenomic discordance, and functional variation. (A) Chloroplast-to-mitochondrion transfers (NUPTs) and nuclear-mitochondrial homology. The *mtcp15* fragment (green) is specific to *C. sinensis* var. *sinensis* cultivars. (B) Chromosomal distribution of nuclear-mitochondrial shared sequences in CSSBY. Phylogenomic conflicts among chloroplast (C), mitochondrial (D), and nuclear genomes (E). Divergence times (red, million years) and bootstrap values (>70% shown) highlight cytonuclear discordance. (F) Presence (black), absence (white), and pseudogenization (gray) of protein-coding, tRNA, and rRNA genes. Copy numbers are indicated for duplicated genes (e.g., *atp9* in CSSBY).

Nuclear-mitochondrial homology spanned 2.32–3.88 Mb across species, with CRE showing the highest nuclear-shared sequences (3,877,865 bp; **Table 1**).

Chromosome 9 of the CSSBY nuclear genome shared 801,420 bp with CRE’s mitogenome, enriched in retrotransposon-derived regions (**Fig. 4B**). Annotation revealed 57–60 mitochondrial genes with nuclear homologs, including 38 protein-coding genes (e.g., *rps3*, *ccmB*) and 21 tRNAs (**Supplementary Table 6**). These findings align with reports of frequent nuclear-mitochondrial DNA exchange in *Arabidopsis* ^32^, but the scale in *Camellia* is unprecedented.

Mitochondrial, chloroplast, and nuclear phylogenies exhibited stark conflicts (**Fig. 4 C-E**). The chloroplast tree (**Fig. 4C**) strongly supported monophyly of *C. sinensis* varieties (bootstrap = 100%), whereas the mitochondrial tree placed *C. impressinervis* (CIM) sister to *C. reticulata* (CRE) with weak support (bootstrap = 76%; **Fig. 4D**). Nuclear phylogeny mirrored chloroplast topology but conflicted with mitochondrial gene trees, particularly in the placement of *C. lanceoleosa* (CL) and *C. taliensis* (CTA) (**Fig. 4E**).

Mitochondrial SNPs and indels further illuminated hybridization signals. CSSLJ43 and CTA, both putative hybrids, showed elevated SNP densities in intergenic regions (1,068 and 766 SNPs, respectively; **Table 1**, **Fig. 2C, Supplementary Table 7**), contrasting with CSSBY’s low polymorphism (605 SNPs). Synonymous-to-nonsynonymous SNP ratios (*dN*/*dS*) averaged 8.7 for protein-coding genes, reflecting strong purifying selection, yet intergenic regions accumulated 4.2× more variants (**Fig. 2C**). This pattern mirrors findings in hybrid *Silene* mitogenomes^33^, suggesting relaxed selection in hybrid genomic backgrounds. Across *Camellia* species, mitochondrial expansion correlates with hybrid features. For example, CRE (788,876 bp) shows 1.2× size reduction compared to CLA, coinciding with retrotransposon-like repeat activation, a pattern associated with hybridization-induced genomic shock.

Mitochondrial gene content was largely conserved, with 65–77 genes per genome (**Table 1**). Protein-coding genes varied minimally (38–46), but tRNA/rRNA counts fluctuated significantly (18–26 tRNAs; 3–6 rRNAs; **Fig. 4F**). Notably, CRE lacked *nad9*, while CLA and CIM independently lost *sdh3* and *mttB*, respectively (**Fig. 4F**). Duplicated genes were rampant: CRE retained two copies of *sdh3*, *sdh4*, *cox3*, and *rps19*, whereas CSSBY maintained single-copy status for all genes (**Fig. 4F**). RNA editing sites numbered 500–527 across species, with *ccmFn* hosting the highest density (34–36 sites; **Supplementary Table 8**). Editing sites were highly conserved: 489/518 (94.4%) sites were shared among all *Camellia* species, exceeding the 87% conservation reported in *Brassica* ^34^.

However, *C. reticulata* (CRE) and *C. lanceoleosa* (CLA) uniquely lacked editing at *nad* and sdh3, respectively, potentially impacting their protein functions (**Supplementary Table 8**). Transcriptomic analysis has identified 496 RNA editing sites in *Camellia lanceoleosa*, which is slightly lower than the results predicted by the software, but is generally reliable (**Supplementary Table 9**).

## Discussion

### Multichromosomal Mitochondrial Architectures: Challenging the Master Circle Paradigm

While nuclear data suggest all investigated *Camellia* species may have hybrid origins^11,12,35^, mitochondrial genomes uniformly capture this history through three hallmarks: short repeat-driven inflation (**Fig. 3C**), rampant HGT (**Fig. 4A, B**), and phylogenomic discordance (**Fig. 4C-E**). Variation in genomic shock intensity, quantified by SNP density (**Table 1**) and repeat profiles, explains lineage-specific differences like CRE’s size reduction and CTA’s multichromosomal architecture.

Our discovery of fragmented, multichromosomal mitogenomes in *C. sinensis* var. *sinensis* cv. Longjing43 (CSSLJ43) and *C. taliensis* (CTA) challenges the long-standing “master circle” model of plant mitochondrial architecture. While multipartite mitochondrial genomes have been reported in *Silene*, *Salvia*, and maize^24,36–38^, their stable maintenance in *Camellia*, a genus not previously associated with such complexity, suggests that fragmented structures may be more widespread in angiosperms than currently recognized. The two-chromosome configuration in CSSLJ43 and CTA was rigorously validated via split-read mapping, ruling out assembly artifacts. These findings align with recent calls to abandon the circular genome assumption in favor of dynamic, network-like models^39^, particularly in lineages with high recombination rates.

The structural divergence between CSSLJ43 and CTA, despite their close phylogenetic relationship, highlights the role of lineage-specific evolutionary pressures. For instance, CTA’s Chr2 (126 kb) is enriched in repeats (>80% of its length), whereas CSSLJ43’s Chr2 (124 kb) harbors functional genes (*nad9*, *rps19*; **Fig. 1B**). This dichotomy mirrors observations in *Silene*, where repeat accumulation correlates with chromosome fragmentation^33^. Our results suggest that repeat-mediated recombination, rather than phylogenetic distance, drives mitochondrial structural diversity in *Camellia*. In addition, frequent interspecific gene flow in *Camellia*^17^ creates a genomic landscape where mitochondrial diversity archives population-level introgression.

### Short Repeats as Drivers of Genome Expansion: A Lineage-Specific Strategy

Contrary to prevailing models emphasizing long repeats (>1 kb) in mitogenome expansion^25,28^, we demonstrate that short repeats (<500 bp) dominate mitochondrial genome inflation in *Camellia*. These repeats account for 92.6–94.9% of all repetitive elements and contribute >70% of total repeat length in most species. Strikingly, the correlation between genome size and total repeat number (*R*² = 0.9729) surpasses values reported in *Cucurbita* ^28^(*R*² = 0.84), suggesting that short repeats are particularly potent drivers of expansion in Ericales.

The exceptional case of *C. taliensis* (CTA), where repeats >1 kb occupy 95.3% of total repeat length, hints at lineage-specific evolutionary trajectories. Species with strong hybridization signatures (e.g., CTA)^40^ may drive repeat accumulation through genomic instability and relax selection against repeat accumulation, allowing long repeats to persist, a phenomenon observed in polyploid *Triticum* species^41^. This duality, short repeats dominating diploids versus long repeats in polyploids, suggests that genome duplication reshapes mitochondrial repeat dynamics, a hypothesis warranting testing in other polyploid-rich clades.

### Horizontal Gene Transfer and Nuclear-Organellar Conflicts

Horizontal gene transfer (HGT) emerged as a pervasive force in *Camellia* mitogenome evolution. Plastid-derived sequences (NUPTs) spanned up to 40 kb (CSSLJ43), exceeding the 15–30 kb range reported in *Brassica*^34^.

The *mtcp15* fragment, shared exclusively among *C. sinensis* var. *sinensis* cultivars, provides the first evidence of lineage-specific chloroplast-to-mitochondrion transfers in *Camellia*. Such events may facilitate mitochondrial adaptation to nuclear-cytoplasmic incompatibilities arising from hybridization, as proposed in *Helianthus*^27^.

Nuclear-mitochondrial homology was equally striking, with *C. reticulata* (CRE) sharing 3.88 Mb with the CSSBY nuclear genome. This scale of nuclear DNA infiltration into mitochondria surpasses findings in *Arabidopsis*^10,32^ and likely reflects *Camellia*’s propensity for hybridization, which may destabilize organellar-nuclear DNA boundaries. Notably, chromosome 9 of the nuclear genome contributed 801 kb to CRE’s mitogenome, a pattern reminiscent of mitochondrial “NUMT hotspots” in humans^42^. These transfers, while typically nonfunctional, could inadvertently provide raw material for mitochondrial innovation.

### Phylogenomic Discordance: Mitochondria as Hybridization Barometers

The pronounced conflict between mitochondrial, chloroplast, and nuclear phylogenies underscores mitochondria’s unique evolutionary trajectory in *Camellia*. Mitochondrial trees placed *C. impressinervis* (CIM) sister to *C. reticulata* (CRE) with weak support (bootstrap = 76%), contradicting chloroplast and nuclear topologies. This discordance likely stems from three factors: 1)Hybridization: Mitochondrial capture during interspecific crosses, as documented in other flowering plants^33^ and suggested by cytonuclear discordance in *Camellia*^14^; 2) Incomplete Lineage Sorting: Mitochondria’s low substitution rate^43^ preserves ancestral polymorphisms erased in faster-evolving nuclear genomes; 3) Differential Selection: Purifying selection on mitochondrial genes (*dN/dS* = 0.12) may slow divergence, whereas nuclear genes experience relaxed selection in hybrids.

The elevated SNP density in CSSLJ43 and CTA, both putative hybrids, mirrors patterns in synthetic *Arabidopsis* hybrids^44^, suggesting that hybridization destabilizes mitochondrial sequence integrity. These findings position mitochondrial genomes as sensitive indicators of historical hybridization, complementing nuclear and chloroplast data in resolving complex evolutionary histories.

### Toward a Mitochondrial-Aware Taxonomy in *Camellia*

Our study demonstrates that mitochondrial structural variants, repeats, HGT events, and chromosomal configurations, provide novel markers for *Camellia* systematics. For instance, the presence of *mtcp15* exclusively in *C. sinensis* var. *sinensis* cultivars offers a molecular synapomorphy to distinguish this group from *C. sinensis* var. *assamica*. Similarly, the two-chromosome structure in CSSLJ43 and CTA could serve as a karyotypic marker for hybrid lineages.

These mitochondrial signatures, when integrated with nuclear and chloroplast data, may resolve long-standing taxonomic ambiguities. For example, *C. taliensis*’s mitochondrial affinity with *C. reticulata* supports its hybrid origin from *C. reticulata* and *C. sinensis* progenitors, as hypothesized by Huang et al. (2014)^12^. This approach mirrors successful efforts in *Pinus*, where mitochondrial rearrangements clarified species boundaries^45^.

## Conclusion

This study establishes *Camellia* as a paradigm for resolving evolutionary paradoxes in flowering plants through comparative mitochondrial genomics. By integrating multi-omics analyses of eight *Camellia* species and their outgroup, we reveal how mitochondrial genome dynamics reflect the interplay of hybridization, repeat-driven expansion, and structural innovation, three forces that have long obscured taxonomic boundaries in this notoriously complex genus. Our findings redefine mitochondrial roles in plant speciation and provide a roadmap for studying cytoplasmic evolution in reticulate systems.

Central to this redefinition is the discovery that short repeats (<100 bp) drive∼92.6–94.9% of mitochondrial genome expansion (*R*² = 0.9729), overturning the long-held view of large repeats as primary drivers. This “micro-repeat dominance” correlates with lineage-specific hybridization events, where introgressed nuclear genomes likely destabilize repeat repair mechanisms. In diploid lineages exhibiting hybrid genomic signatures (e.g., CRE with retrotransposon-like repeats; CTA with fragmented chromosomes), mitochondrial expansion reflects lineage-specific responses to introgression-driven genomic shock, rather than polyploidy.

The multichromosomal mitochondrial architectures in *C. sinensis* var. *sinensis* LJ43 and *C. taliensis* (CTA) exemplify structural plasticity as an adaptive strategy. Chromosome fission events, validated by split-read mapping and PCR, create functionally autonomous genomic modules enriched in energy metabolism genes (*nad9*, *rps19*). These modules exhibit reduced recombination rates at repeat-rich junctions, potentially buffering against hybrid breakdown.

Phylogenetically, fragmented mitochondrial genomes correlate with recent radiation events in *Camellia*, implying that structural modularity facilitates rapid diversification in taxonomically contentious groups.

Horizontal gene transfer (HGT) emerges as both a driver and recorder of reticulate evolution. Plastid-derived sequences (up to 25.5% of mitochondrial content) show lineage-specific patterns: the *mtcp15*transfer event unites three *C. sinensis* varieties, postdating their divergence from *C. assamica*. Nuclear-mitochondrial homology hotspots on chromosome 9 (801,420 bp in CRE) align with regions of ancestral introgression inferred from nuclear phylogenies, suggesting HGT mitigates cytonuclear conflict during hybridization. These transfers create a mosaic genomic landscape where mitochondrial diversity mirrors the genus’s complex breeding history.

The phylogenetic discordance between mitochondrial, chloroplast, and nuclear trees resolves long-standing taxonomic ambiguities. Mitochondrial data clarify the sister relationship between *C. impressinervis* (CIM) and *C. reticulata* (CRE), a grouping obscured by chloroplast introgression, while exposing hybridization signatures in CSSLJ43 through elevated intergenic SNP densities. Such conflicts underscore mitochondria as “evolutionary recorders” of historical gene flow, providing complementary signals to nuclear data for reconstructing reticulate phylogenies.

By bridging mitochondrial genomics with taxonomy in a hybridization-prone system, this work transforms *Camellia* from a taxonomic enigma into a model for studying cytoplasmic evolution. The micro-repeat expansion mechanism and chromosomal modularity theory advance fundamental understanding of organellar genome plasticity, with implications for resolving species complexes in *Rhododendron*, *Quercus*, and other hybrid-rich genera. Future efforts should explore how mitochondrial-nuclear coevolution shapes reproductive isolation and whether repeat-driven genome expansion imposes metabolic costs affecting ecological adaptation. This framework positions mitochondrial genomics as an indispensable tool for deciphering the tangled evolutionary histories that characterize much of angiosperm diversity.

In summary, by unraveling the interplay of repeats, HGT, and hybridization in shaping *Camellia* mitogenomes, this study redefines principles of plant mitochondrial evolution. Our findings challenge the universality of long-repeat-driven expansion, expose mitochondria as archives of hybridization history, and provide tools to navigate taxonomic complexities in radiations shaped by introgression.

## Materials and Methods

### Plant Materials and DNA Sequencing

Fresh leaves from eight *Camellia* species (*C. sinensis* var. *sinensis* cv. Biyun (CSSBY), *C. sinensis* var. *sinensis* cv. Fuding (CSSFD), *C. sinensis* var. *assamica* cv. Yunkang (CSAYK10), *C. lanceoleosa* (CLA), *C. impressinervis* (CIM), *C. taliensis* (CTA), and *C. reticulata* (CRE)) were collected from Yunnan Province, China. Voucher specimens were deposited at the Herbarium of Hainan University (voucher codes: HUTB-2025-001 to HUTB-2025-008). Total genomic DNA was extracted using an improved CTAB protocol^46^, with RNA removed by RNase A treatment (Thermo Fisher, USA). For PacBio sequencing, 20 µg of high-molecular-weight DNA (HMW DNA) per sample was sheared to 20 kb using a Megaruptor 3 system (Diagenode, Belgium) and processed with the SMRTbell Express Template Prep Kit 2.0 (Pacific Biosciences, USA). Libraries were sequenced on the PacBio Sequel II platform (Menlo Park, CA) with 30-hour movie times, generating ∼130× coverage per sample (**Supplementary Table 10)**. Illumina paired-end libraries (350 bp insert size) were constructed using the NEBNext Ultra II DNA Library Prep Kit (NEB, USA) and sequenced on the NovaSeq 6000 platform (Illumina, USA) to generate 150 bp reads.

### Mitochondrial Genome Assembly and Annotation

Mitochondrial reads were extracted by mapping PacBio subreads to the reference CSAYK10 mitogenome^47^ (GenBank: MK573092) using BWA-MEM^48^ v0.7.17. *De novo* assembly was performed with CANU^49^ v1.8 under default parameters, followed by three rounds of error correction using Pilon^50^ v1.23 with Illumina data. Contigs were validated by the average depth of reads mapped to each contig.

The mitochondrial genome was annotated using Mitofy^28^ for protein-coding genes (PCGs), tRNAscan-SE^51^ v2.0 for tRNAs, and RNAmmer^52^ v1.2 for rRNAs.

Chloroplast-derived sequences were identified by BLASTN against the CSSBY chloroplast genome (GenBank: MN539999), with hits >100 bp and E-value <1×10^−5^ retained. Genome maps were visualized with OGDRAW^53^ v1.3.1 and manually refined in Adobe Illustrator.

### Repeat Sequence Analysis

Simple sequence repeats (SSRs) were detected using MISA^54^ v2.1 with thresholds of 8, 4, 4, 3, 3, and 3 repeats for mono-to hexanucleotides. Tandem repeats (TRs) were identified with Tandem Repeats Finder^55^ v4.07 using default settings. Long repeats (>50 bp) were detected with REPuter^56^ under parameters of minimal repeat length 50 bp, Hamming distance 3, and E-value <1×10^−5^. Repeat content was classified by size: <100 bp, 100–500 bp, 500–1,000 bp, and >1,000 bp.

### Genome Structural Variants Detection and RNA Editing Site Prediction

SNPs and InDels were identified by aligning genome to the CSSBY mitochondrial genome using MUMmerv3.1^57^. The number of SNPs and InDels were identified using BCFtools v1.10^58^. Circos v0.69-6^59^ was used to display the detected mt genome structural variants. The functional impact of SNPs and InDels was annotated using SnpEff^60^ v4.3t. RNA editing sites in PCGs were predicted using the PREP-Mt server^61^ with a cutoff value of 0.6. Only C-to-U conversions supported by alignments to *Arabidopsis thaliana* mitochondrial transcripts (TAIR10) were retained. The RNA-seq data of the root, stem and leaf samples (SRR15524230, SRR15524234 and SRR15524236) were downloaded from NCBI SRA database. The cleaned reads from the three tissues were mapped to the CLA mitogenome by bowtie2 v2.4.5^62^ with mismatch = 7. These mapped reads were extracted by samtools v1.11^58^ with the parameter “flag = 16” to separate those mapped to the forward and reverse strands. Finally, RNA editing sites from both strands were separately called by REDItools v2^62^ with the following cutoffs: coverage ≥ 5, frequency ≥ 0.1, and *P* ≤ 0.05.

### Horizontal Gene Transfer (HGT) Detection

Nuclear-mitochondrial homologous regions were identified by BLASTN^63^ v2.8.1+ searches against the CSSBY nuclear genome^14^ with thresholds of >100 bp alignment length and E-value <1×10^−5^. Plastid-derived sequences (NUPTs) were detected similarly using the CSSBY chloroplast genome as a reference. Transferred regions were visualized with Circos^59^ v0.69.

### Phylogenetic and Synteny Analyses

Phylogenetic trees were reconstructed using 1) Chloroplast genomes (**Supplementary Fig. 1B**): 80 conserved genes aligned with MAFFT^64^ v7.475, trimmed by trimAl^65^ v1.4, and analyzed with RAxML^66^ v8.2.12 under the GTR+GAMMA model; 2) Mitochondrial genomes: 26 conserved PCGs concatenated and analyzed as above; and 3) Nuclear orthologs: Single-copy genes identified by OrthoFinder^67^ v2.5.2, aligned, and processed with IQ-TREE^68^ v2.1.2 under the best-fit model (TESTNEW). To generate a linear comparison of the gene neighborhood around the mt genomes, translated BLAST v2.8.1+^63^ was performed with maximum E-value set to 1×10^-5^ followed by map generation with Easyfig v2.2.3^69^. Synteny analysis was performed using MUMmer^57^ v3.1 with default parameters, and collinear blocks were visualized with Circos^59^.

## Statistical Analyses

Correlations between genome size and repeat content were assessed using phylogenetic generalized least squares (PGLS) in R v4.2.0 with the *caper* package. Synonymous (*dS*) and nonsynonymous (*dN*) substitution rates were calculated with KaKs_Calculator^70^ v2.0.

## Acknowledgments

This work was supported by Hainan Provincial Science and Technology Talent Innovation Project “Rolling Support” Project(KJRC2025B05)and a startup grant of Hainan University (RZ2100006631) (to Gao LZ).

## Author contributions

L.G. conceived and supervised the research. F. Z. performed the data analysis. L.G. and F. Z. interpreted the results, drafted the manuscript, and revised the manuscript. All authors reviewed the results and approved the final version of the manuscript.

## Data availability

The raw sequence reads, genome assembly, and annotation data generated in this study have been deposited in the China National Center for Bioinformation under accession number C_AA113472.1-C_AA113482.1. The raw sequence data of *Camellia sinensis* var. *sinensis* cv. Longjing43 (CSSLJ43) are available at NCBI under Bioproject accession SRP155011. The raw sequence data of *Actinidia eriantha* (AER) are available at NCBI under Bioproject Accession ERP123025. The mitochondrial reference genome data of CSSYK10 are available at NCBI GenBank accession MK574877.

## Supplementary Figures and Tables

**Supplementary Fig. 1 | Mitochondrial genome validation and chloroplast genome structure of *Camellia* species.** (A) Evaluation of the mitochondrial genome assemblies of *Camellia* species. Calculation of depth by mapping reads to the final assembled genomes; (B)The chloroplast genome of *Camellia* species. Genes outside and inside the circle are transcribed clockwise and counterclockwise, respectively. Functional groups are color-coded. Inner ring shows GC content (gray gradient) and AT content (light gray). Genome sizes range from 154,677 to 163,809 bp.

**Supplementary Fig. 2 | Mitochondrial gene order conservation across *Camellia* species.** Lines connect orthologous genes between CSSBY (reference) and other mitogenomes. Non-collinear blocks indicate structural rearrangements. Gene categories: CDS (green), tRNA (blue), rRNA (orange). Species arranged by nuclear phylogeny (**Fig. 5C**). Assembly gaps (<5% of genomes) are omitted for clarity.

**Supplementary Fig. 3 | Regression analyses of mitochondrial genome size versus repeat content.** (A-J) Subgroup analyses for repeat size categories: <100 bp (A-B), 100-500 bp (C-D), 500-1,000 bp (E-F), >1,000 bp (G-H), ≤500 bp (I-J). (K) Repeats >100 bp. PGLS regression lines (solid) with 95% confidence intervals (dashed). Species labels: CSSBY (circle), CSSFD (square), CSSLJ43 (triangle), CSAYK10 (diamond), CT (pentagon), CL (hexagon), CRD (cross), CI (star).

**Supplementary Table 1 | Comparisons of base transition and transversion between *CSSBY* and the seven Theaceae mitochondrial genomes.**

**Supplementary Table 2 | Simple sequence repeat (SSR) profiles.** Counts of mono-to hexanucleotide SSRs across species. Dimeric repeats dominate (257-348 loci), while hexamers are rare (0-4 loci). Detection thresholds: 8 (mono), 4 (di/tri), 3 (tetra/penta/hexa) repeats.

**Supplementary Table 3 | Repeat content classification by size.** Genome size, repeat counts, and cumulative lengths for four size categories. Short repeats (<100 bp) constitute 92.6-94.9% of total repeats in diploids.

**Supplementary Table 4 | Tandem repeat (TR) annotations.** TR coordinates, period sizes, copy numbers, and sequence alignments for all species. TRs ≥6 bp with ≥70% identity were identified using Tandem Repeats Finder v4.07 (Benson, 1999).

**Supplementary Table 5 | Long repeat (>50 bp) characteristics.** Repeat counts, total lengths, and maximal repeat sizes. *C. taliensis* (CT) exhibits exceptional repeat expansion (>50 kb repeats occupy 48.5% of its genome).

**Supplementary Table 6 | Nuclear-mitochondrial gene homology.** Presence/absence of mitochondrial gene homologs in nuclear genomes. “+” indicates homology (BLASTN, E-value <1×10⁻⁵). Nuclear chromosome 9 is a hotspot for mitochondrial DNA transfers.

**Supplementary Table 7 | SNPs and InDels detected by comparing *CSSBY*** mitochondrial genome to the seven other Theaceae mitochondrial genomes.

**Supplementary Table 8 | RNA editing site distribution in mitochondrial genes.** C-to-U editing sites predicted by PREP-Mt (Mower, 2005) with a cutoff score of 0.6. *C. reticulata* (CRE) lacks editing at *nad9*-32 and *ccmFn*-15. Conservation rates calculated relative to *Arabidopsis thaliana*.

**Supplementary Table 9 | Transcriptome-based validation of RNA editing sites in** C. lanceoleosa.

**Supplementary Table 10 | PacBio sequencing statistics and assembly metrics.** Subread counts, N50 lengths, sequencing depth, and assembly validation details for eight *Camellia* mitogenomes and *A. eriantha*.

