## Supplementary figures and images for "Short repeats drive mitochondrial genome expansion and record hybridization in *Camellia*"

### Supplementary Figure 1.pdf

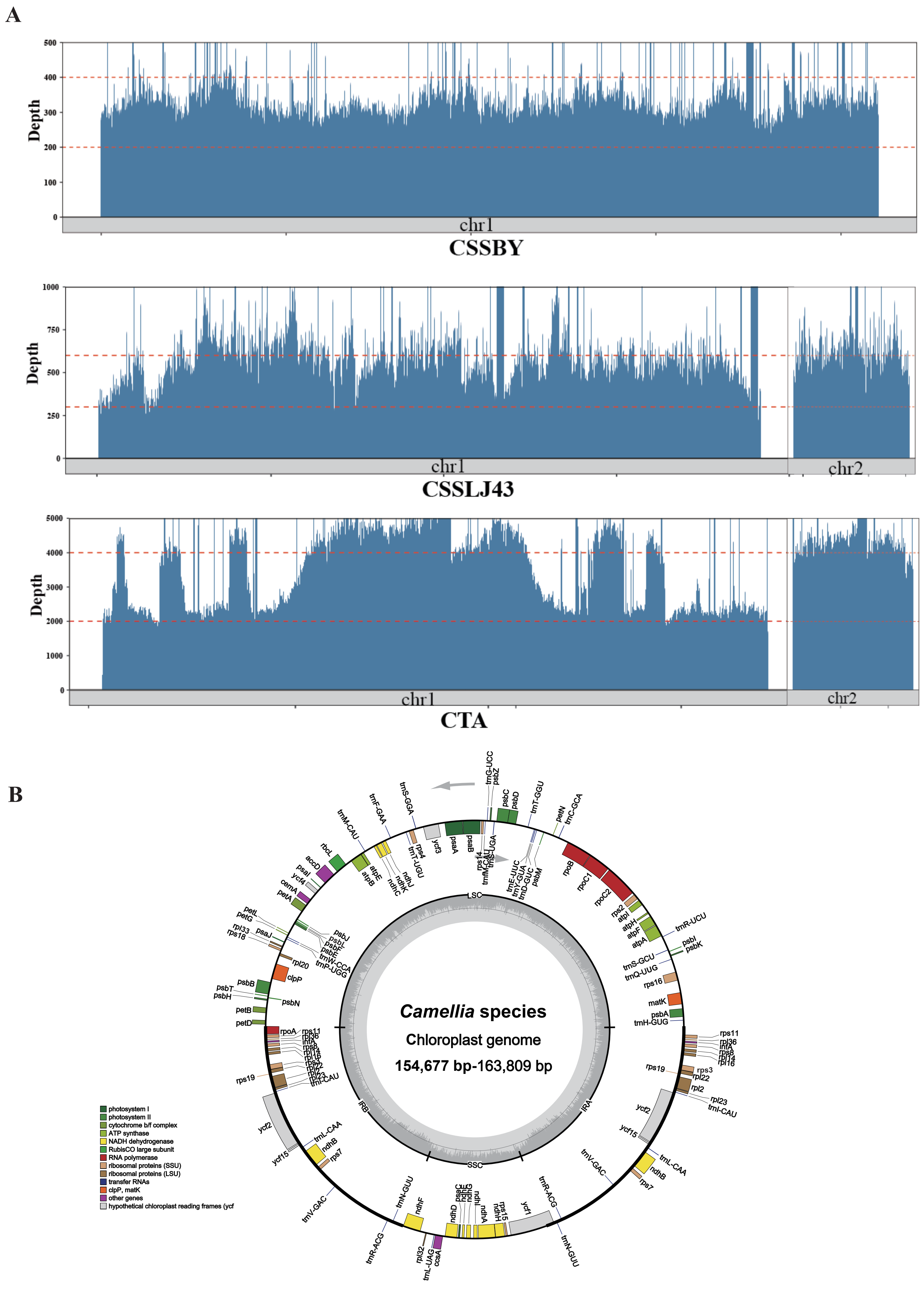

### Supplementary Figure 2.pdf

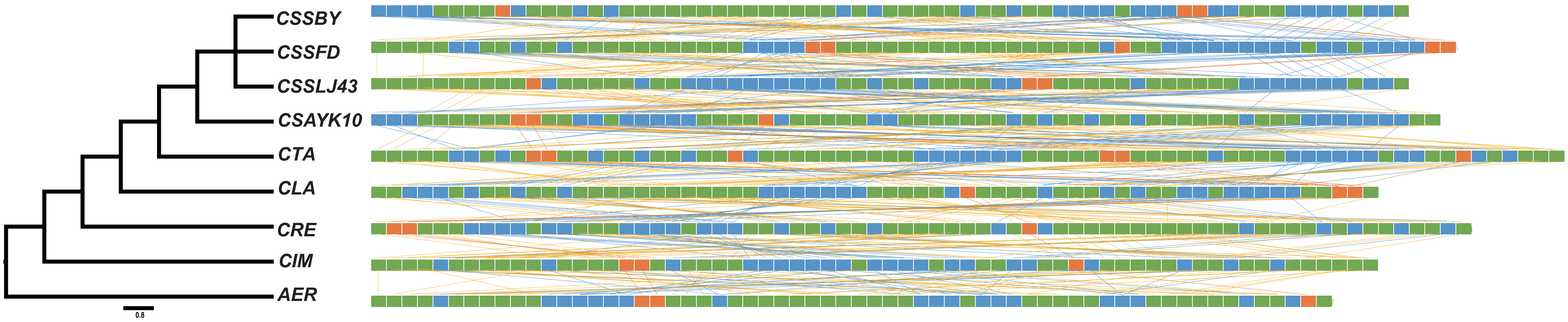

### Supplementary Figure 3.pdf

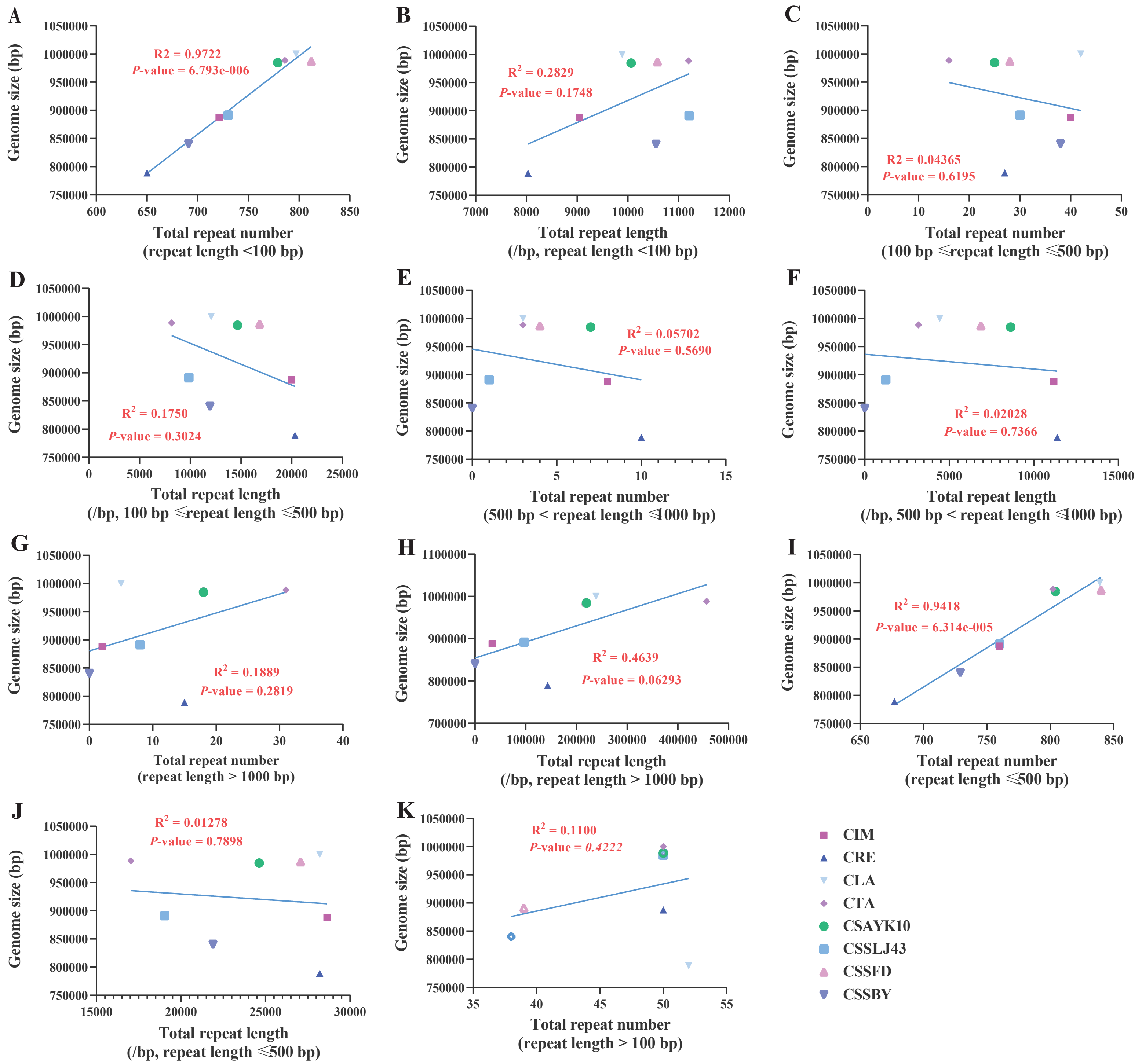
